# Phosphatases are predicted to govern prolactin-mediated JAK-STAT signaling in pancreatic beta cells

**DOI:** 10.1101/2021.10.14.464474

**Authors:** Ariella Simoni, Holly Huber, Senta K. Georgia, Stacey D. Finley

**Author notes:** Corresponding author Stacey D. Finley, 1042 Downey Way, DRB 140, Los Angeles, CA 90089.

## Abstract

Patients with diabetes are unable to produce a sufficient amount of insulin to properly regulate their blood-glucose levels. One potential method of treating diabetes is to increase the number of insulin-secreting beta cells in the pancreas to enhance insulin secretion. It is known that during pregnancy, pancreatic beta cells proliferate in response to the pregnancy hormone, prolactin. Leveraging this proliferative response to prolactin may be a strategy to restore endogenous insulin production for patients with diabetes. To investigate this potential treatment, we previously developed a computational model to represent the prolactin-mediated JAK-STAT signaling pathway in pancreatic beta cells. However, this model does not account for variability in protein expression that naturally occurs between cells. Here, we applied the model to understand how heterogeneity affects the dynamics of JAK-STAT signaling. We simulated a sample of 10,000 heterogeneous cells with varying initial protein concentrations responding to prolactin stimulation. We used partial least squares regression to analyze the significance and role of each of the varied protein concentrations in producing the response of the cell. Our regression models predict that the concentrations of the cytosolic and nuclear phosphatases strongly influence the response of the cell. The model also predicts that increasing prolactin receptor strengthens negative feedback mediated by the inhibitor SOCS. These findings reveal biological targets that can potentially be used to modulate the proliferation of pancreatic beta cells to enhance insulin secretion and beta cell regeneration in the context of diabetes.

## Introduction

The pancreas is the primary organ able to mediate glucose homeostasis, releasing various hormones, including insulin, to maintain blood glucose levels. Diabetes is a disease in which patients are unable to produce enough insulin to properly maintain blood-glucose homeostasis. In the case of Type 1 diabetes, insulin insufficiency occurs because of autoimmune destruction of insulin secreting beta cells; in the case of Type 2 diabetes, prolonged insulin resistance leads to beta cell dysfunction and insufficient insulin production. It is possible that increasing the number of functional insulin-secreting beta cells in the pancreas could be a means to alleviating diabetes.

Interestingly, in certain physiological conditions, pancreatic beta cells expand to respond to increased metabolic demand for insulin. For example, beta cells adapt to pregnancy-associated insulin resistance by increasing beta cell mass and insulin secretion [1]. Expression of lactogenic hormones, such as prolactin (PRL), promote beta cell proliferation in early pregnancy [2,3]. Prolactin receptor (PRLR) has been identified as the receptor responsible for integrating lactogenic signaling in beta cell mass during pregnancy [3–5]. This pathway could be harnessed to regenerate beta cell mass as a novel therapy to treat insulin-dependent diabetes.

In beta cells, PRL-mediated activation of the PRLR activates the JAK-STAT signaling cascade, which results in the activation of Signal Transducer and Activator of Transcription 5 (STAT5) and mediates downstream signaling [6,7]. Specifically, STAT5 is a transcription factor for cell cycle proteins, such as D-type cyclins, and anti-apoptotic proteins, such as BcL-xL [8,9]. Together, the signaling cascade initiated by these proteins promotes beta cell proliferation. Yet, beta cells do not all proliferate simultaneously; this suggests there is a heterogeneity that naturally occurs in a population of beta cells [10] that may be governed by intracellular signaling.

The JAK-STAT signaling pathway that is responsible for beta cell proliferation is complex, containing many species and reactions. Computational models are useful in studying such complex signaling pathways. In particular, models allow us to investigate the effects of modulating aspects of the pathway, which would otherwise be difficult to do. Mathematical modeling has been applied to study different aspects of pancreatic beta cell signaling [11] and metabolism [12], providing insight into mechanisms underlying cellular behavior. We recently developed a novel computational model to represent the JAK-STAT signaling cascade [13]. The model was fitted to match experimental data of pancreatic beta cells from the INS-1 cell line treated with PRL. However, the effects of heterogeneity on this model have yet to be analyzed. Therefore, in this study, we incorporate heterogeneity into the model in order to further understand the mechanisms of the JAK-STAT signaling pathway, and to identify potential strategies to modulate the population-level response to the signaling cascade.

We used the computational model to simulate a population of 10,000 cells responding to PRL treatment. We studied how heterogeneity impacts this model by varying the initial protein concentrations of each of the 10,000 cells prior to simulation. In this work, we assume the response of each cell was given by the levels of phosphorylated STAT5, the transcription factor as it mediates production of PRLR and the anti-apoptotic protein, BcL-xL [8,14,15]. We used a data-driven approach of partial least squares regression analysis to relate the initial protein concentrations in the cells to the resulting cellular responses. Using PLSR analysis, we identified the relative importance of each of the proteins in producing the response of the cell at a single time point. Additionally, we analyzed the overall shapes and features of the time course response. Overall, our combination of mechanistic and data-driven modeling uncovers the roles of intracellular protein concentrations in affecting specific aspects of the signaling response.

## Methods

### Modeling framework

This study utilized an ordinary differential equation (ODE) model of the JAK2-STAT5 signaling pathway in pancreatic beta cells that we previously developed by Mortlock et al [13]. (**Figure 1**). Signal transduction begins when PRL binds to PRLR, which is bound to the JAK2 kinase. PRL binding initiates dimerization of the PRLR-JAK2 complex and activation of JAK2. The activated PRLR-JAK2 complex mediates the phosphorylation of the transcription factor STAT5. Phosphorylated STAT5 (pSTAT5) dimerizes, and upon dimerization, translocates to the nucleus to promote transcription of its target genes. These target genes transcribe mRNA encoding the PRLR-JAK2 complex, anti-apoptotic protein BcL-xL, and suppressor of cytokine signaling (SOCS) protein. Finally, these three mRNA transcripts are translated into the respective proteins. In addition to inhibition by SOCS, signaling is attenuated by phosphatases in the cytosol and the nucleus, as well as ligand-induced receptor internalization. The rates of the biochemical reactions in this signaling cascade are represented using mass-action and Michaelis-Menten kinetics.

**Figure 1.**
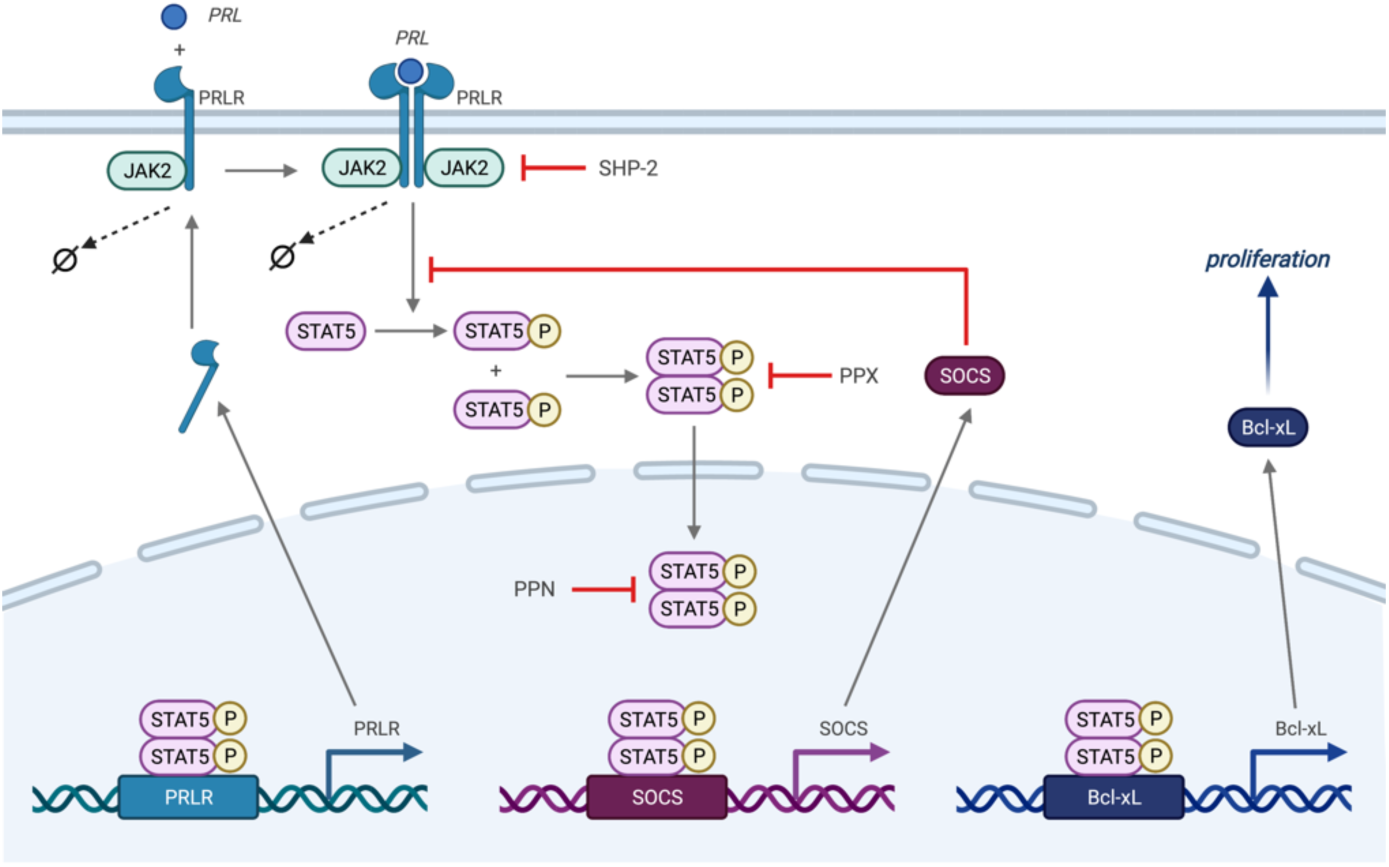
Mechanistic model schematic. Illustration of the signaling cascade modeled in this work. PRL binds to the PRLR-JAK2 complex, leading to receptor dimerization and JAK activation. The activated complex promotes phosphorylation of STAT5, which dimerizes and can translocate to the nucleus. Nuclear STAT5 promotes transcription and translation of PRLR, BcL-xL, and SOCS. Feedback from these proteins and the effect of phosphatases are indicated.

### Parameter estimation

Bayesian inference was used to estimate the kinetic parameters. This approach approximates the conditional probability of the model parameters given experimental data, or the posterior distribution. The posterior distribution is proportional to the likelihood distribution (the conditional probability of the experimental data given the model parameters) multiplied by the prior distribution, or prior probability of the parameters. This relationship is specified by Bayes Theorem:

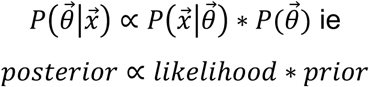

Where:

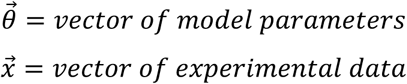

We implement Bayesian parameter estimation using the Metropolis-Hastings algorithm, and most of our protocol mirrors the approach used in previous work [16,17]. Supplementary Table 1 includes all parameters used to define the distribution of the likelihood, prior, and proposal. All are consistent with Mortlock et al., except for the variance of the proposal distribution, which was tuned to achieve an acceptance rate between 0.15 and 0.5. An acceptance rate within this range has been shown to be at least 80% efficient [18]. Additionally, we improve our ability to monitor the Metropolis-Hastings algorithm’s convergence to the posterior distribution. In contrast to the original posterior approximation, we initiated three independent sample chains and monitored convergence via quantitative and qualitative diagnostics. These diagnostics included the Gelman-Rubin (G-R) method (improved upon by Brooks in 1998), autocorrelation plots, trace plots, and histogram plots for each marginal posterior distribution [19]. Posterior sampling ended when the G-R method dropped below 1.2, a rule-of-thumb threshold for algorithm convergence [20]. More detail about the G-R method is provided in the supplemental material.

We sampled our chains from the prior distribution used in Mortlock et al. The log-normal shape and variance of 2 for this distribution is derived from an aggregated dataset of kcat parameters across cell types and intracellular cell pathways [21]. Because our model’s posterior distribution describes only one cell type and one pathway, this distribution was assumed to be over-dispersed with respect to our posterior distribution. Importantly, since we initiate chains from an over-dispersed distribution and improved upon our ability to monitor convergence of the Metropolis-Hastings algorithm, we can be more confident that this parameterization represents a global, rather than local, optima for in silico experimentation.

Each chain had 3 × 10^5^ samples of the posterior distribution after convergence was concluded. The ODE model was parameterized with each of the 9 × 10^5^ posterior samples, and the sample producing the lowest squared difference between model predictions and experimental data was chosen for model simulations.

### Simulating heterogeneity

In this study, we investigated how cell-to-cell heterogeneity affected the outcome of the model. To do this, we chose to vary initial species concentrations between cell simulations. The model defines 56 unique species; however, most of these species’ concentrations begin as zero at the start of the simulation or have initial concentrations that are calculated based on other constants in the model. Therefore, we chose to vary the species that have a nonzero initial concentration and whose initial concentrations are not calculated based on other constants. In particular, we varied the initial concentrations of the prolactin receptor associated with JAK2 (RJ), SH2 domain-containing tyrosine phosphatase 2 (SHP2), cytosolic phosphatase (PPX), and nuclear phosphatase (PPN). We simulated a population of 10,000 cells, and in each cell, we independently selected initial concentrations of RJ, SHP2, PPX, and PPN. Using MATLAB, we sampled from a log-uniform distribution that ranged 10-fold above and 10-fold below the values predicted by the parameter estimation for each of the four species. We chose this wide sampling range to sufficiently encompass the variability that naturally occurs between cells and capture cell response to extreme deviations from the mean.

### Characterizing cellular response

We simulated each of the 10,000 cells, producing data describing the response over a six-hour time course. The cellular response was characterized by four separate quantities: the nuclear-to-cytosolic ratio of pSTAT5A (*ratio*_A_), the nuclear-to-cytosolic ratio of pSTAT5B (*ratio*_B_), the concentration of pSTAT5A relative to all forms of STAT5A (*relative*_A_), and the concentration of pSTAT5B relative to all forms of STAT5B (*relative*_B_). We chose to measure the activity of the transcription factors to characterize the cell response as opposed to measuring the downstream production of anti-apoptotic protein BcL-xL, since the activity of the transcription factors provide more insight into the upstream activity of the cell, and STAT5 is an important regulator of other downstream signaling species.

### Analyzing cellular response

The time courses of each of the four responses for all 10,000 cells were analyzed. We used MATLAB to identify the time courses that followed a specific shape that qualitatively matched the response observed in experimental data: the response first rises to a peak, then falls to a minimum, and finally increases again. For a time course to be considered this “peak-minimum-recovery” shape, the time course must have only one local maximum and one local minimum, with the local maximum occurring first. Additionally, the slope between the local minimum and the six-hour time point must be positive. Of the time courses that met these requirements, we recorded six features from each time course (**Figure 2**): the height of the peak, the height of the minimum, the time of the peak, the time of the minimum, the absolute value of slope from the peak to the minimum, and the slope from the minimum to the six-hour time point. The absolute value of the slope from the peak to the minimum was used since this feature is, by definition, negative in value, and converting it to a positive value allowed us to take the log of this value, which was necessary for our analysis.

**Figure 2.**
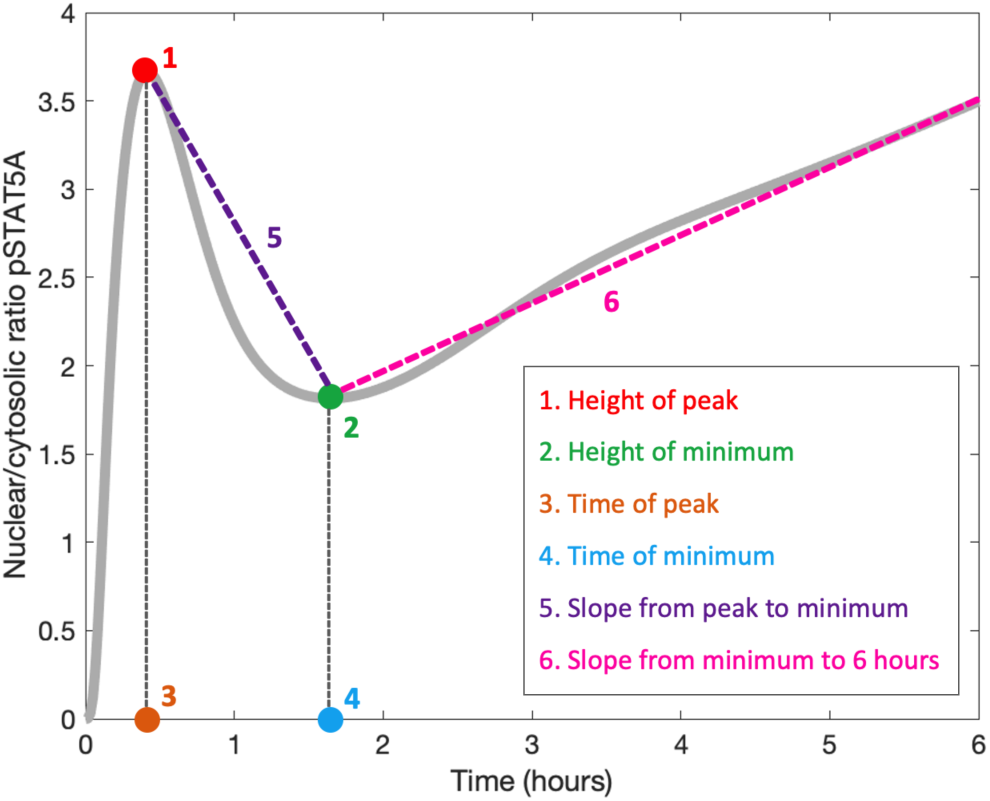
Features of the signaling dynamics investigated. We examined six response features for the STAT5 time course: (1) height of peak, (2) height of minimum, (3) time of peak, (4) time of minimum, (5) absolute value of slope from peak to minimum, and (6) slope from minimum to six hours.

### PLSR analysis

We created partial least squares regression (PLSR) models to approximate the cell response as a function of the initial species concentrations. While a mechanistic model predicts the outputs by simulating individual reactions, a PLSR model predicts the outputs through a linear relationship between the inputs and the outputs, which is much less computationally intensive. This is accomplished by mapping the inputs and outputs into a new space of fewer dimensions, referred to as principal components [22]. In this study, the inputs were the initial species’ concentrations sampled for a given cell, and the outputs were the cell responses predicted by the mechanistic model.

We created PLSR models to analyze the six-hour timepoints of each of the four cell responses (*ratio*_A_, *ratio*_B_, *relative*_A_, *relative*_B_). We made separate PLSR models for each of these outputs, totaling four models. Additionally, we created PLSR models to analyze the six features extracted from the time courses. We made separate PLSR models for each of the six features, for each of the four responses, totaling an additional 24 models. In total, our study included 28 separate PLSR models, each with just one output. With PLSR, it is possible for one model to include multiple outputs; however, we chose to make a separate model for each output since PLSR analysis will only tell you how the inputs affect the overall combined outputs, rather than how the inputs affect each of the outputs separately. Our goal was to uncover how each input influenced each separate output, so we chose to create separate models for each output so we could have this analysis.

We used the data generated by the 10,000 simulations of the mechanistic model to train and test the PLSR models. We trained each model on 90% of the data and tested the models on the remaining 10% of the data. We used the R^2^ value as a metric to quantify the goodness of fit for the testing data.

A useful feature of PLSR modeling is that it calculates the relative contribution of each input in producing the principal components, defined as the weight of the input. The sign of the weight corresponds to whether the input has a positive or negative contribution to each principal component. This allows us to identify the relative importance of each input, as well as the direction in which the input influences the outputs. To identify which inputs are particularly significant in producing the output, we calculate the variable importance of projection (VIP) score of each input, which is determined based on the weight of the input in each component combined with the variance of the output explained by the component. Generally, inputs with a VIP score greater than or equal to one are considered particularly influential in producing the output [23]. In our analysis, we examined the weights and VIP scores to identity which species are influential in producing the cell response and time course features, and in which direction the inputs influence them.

The full model is available at https://github.com/AriellaS/PancreaticBetaCellModeling.

## Results

### Heterogeneity in the dynamic signaling responses

The simulated population of 10,000 cells exhibited a range of time course shapes among the four response quantities following PRL stimulation (**Figure 3**). For each of the four response definitions, many of the time courses followed a general pattern where the response rose to a peak, then fell to a minimum, and finally recovered. This peak-minimum-recovery shape was consistent with the trends observed in the experimental data that the mechanistic model was fitted to. However, some of the time courses did not follow this pattern, with some cells not recovering after the falling phase or some increasing monotonically throughout the entire six hours.

**Figure 3.**
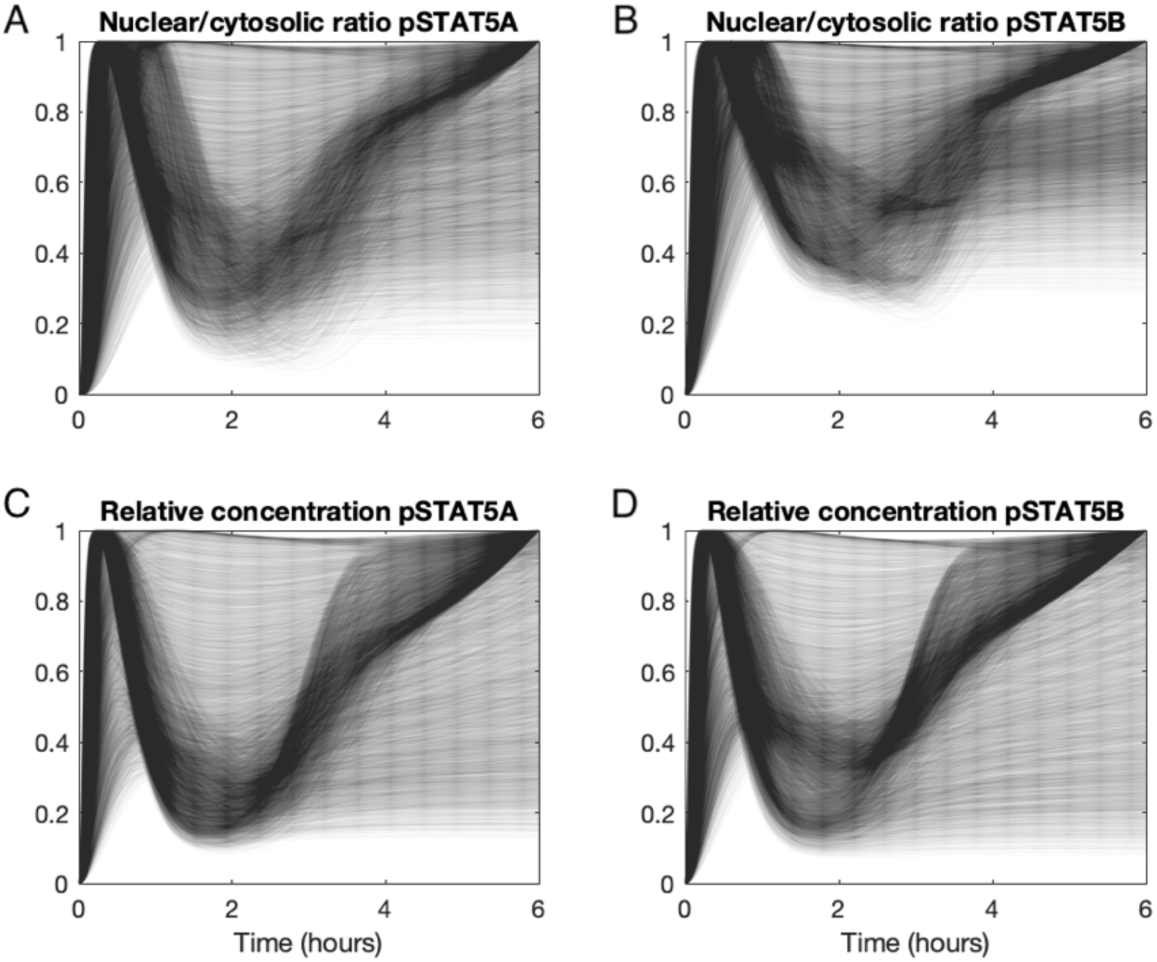
Normalized time courses of cell responses. Responses of each of the 10,000 cells plotted over six hours. Each translucent line represents the response of one cell for (A) Nuclear to cytosolic ratio of pSTAT5A (“*ratio*_B_”), (B) Nuclear to cytosolic ratio of pSTAT5B (“*ratio*_B_”), (C) Relative concentration of pSTAT5A (“*relative*_A_”), and (D) Relative concentration of pSTAT5B (“*relative*_B_”). Each curve is normalized to its respective maximum.

Nearly all the cells followed the peak-minimum-recovery time course shape among the four response quantities. Specifically, 9,574 of the 10,000 simulated cells exhibited the typical peak-minimum-recovery shape under the *ratio*_A_ response definition, with the remaining 426 lacking one or more of the features necessary to fall into that shape. Only the cells that exhibited the peak-minimum-recovery shape for the *ratio*_A_ definition also exhibited the peak-minimum recovery shape for the *ratio*_B_ response definition. 9,570 of the 10,000 cells exhibited the peak-minimum-recovery shape under the *relative*_A_ response definition. These are the same cells that followed the peak-minimum-recovery shape for *ratio*_A_ and *ratio*_B_. 9,569 cells exhibited the peak-minimum-recovery shape under the *relative*_B_ response definition. Only the cells that followed the peak-minimum-recovery shape for *relative*_A_ are grouped into this classification.

### Heterogeneity in the signaling response at six hours

The population of 10,000 cells displayed a wide range of outcomes at the six-hour time point. We first focused on the cell’s response at the six-hour time point in the dynamic response (**Figure 4**). Six hours is an intermediate time point that links short-term signaling of STAT5 phosphorylation and translocation to the longer timescale of the Bcl-xL protein levels (18-24 hours) that influences beta cell proliferation. By focusing on the response at six hours, we can infer how short-term dynamics of STAT5 influence protein production.

**Figure 4.**
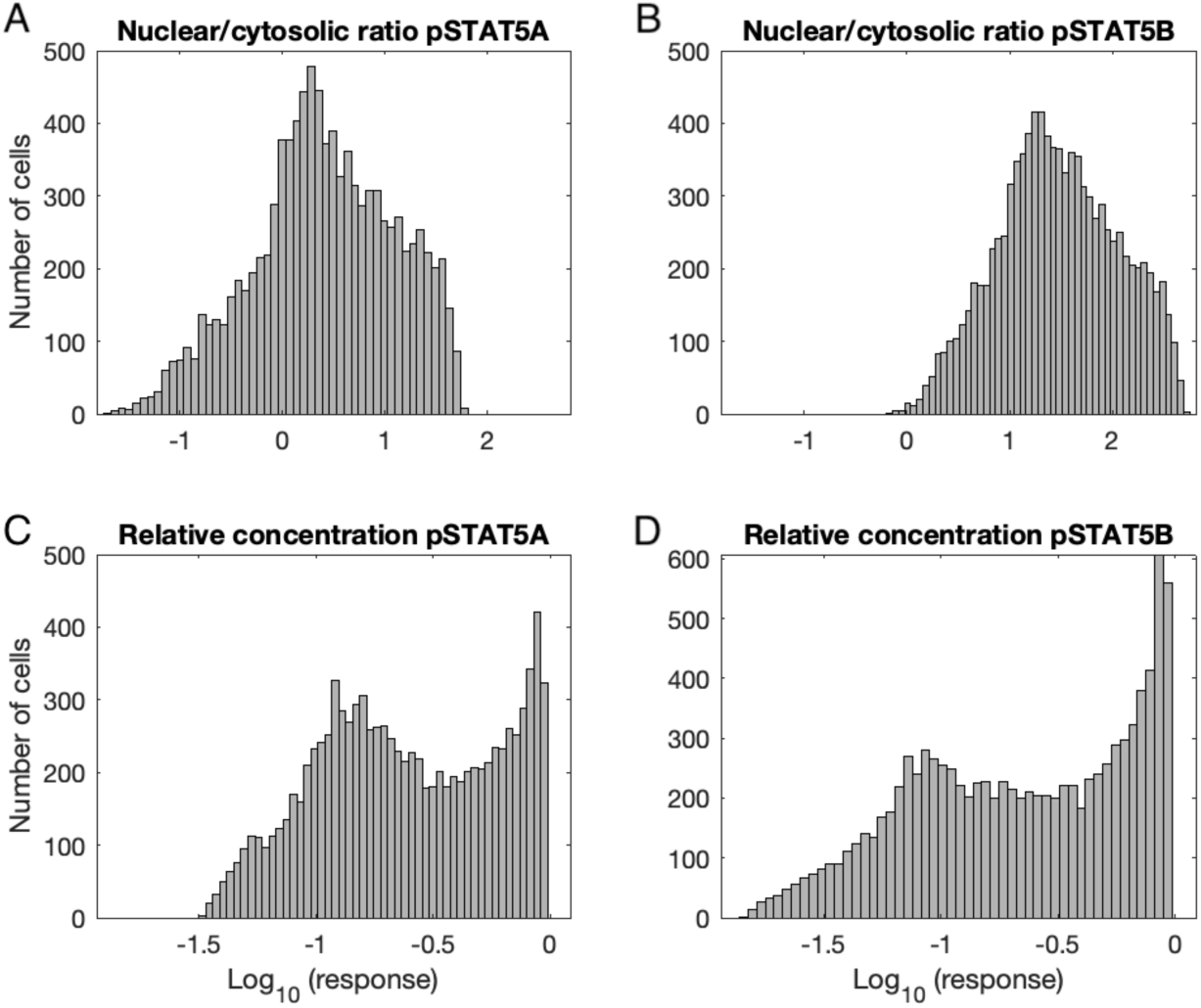
Distributions of cell responses. Distributions of (A) *ratio*_A_, (B) *ratio*_B_, (C) *relative*_A_, and (D) *relative*_B_ at the six-hour time point for the 10,000 simulated cells, shown on a log scale.

The *ratio*_A_ and *ratio*_B_ responses at the six-hour time point exhibited approximately lognormal distributions (**Figure 4A, B**). The median of six-hour *ratio*_A_ was 2.5, while the median of six-hour *ratio*_B_ was 28.1. Thus, the model predicts that *ratio*_B_ tended to be approximately 10-fold greater than *ratio*_A_ at the six-hour time point. The *relative*_A_ and *relative*_B_ responses at the six-hour time point exhibited somewhat bimodal distributions (**Figure 4C, D**). The median of six-hour *relative*_B_ and *relative*_B_ were both 0.23, indicating similar phosphorylation levels of the two proteins.

### Effects of initial protein concentrations on signaling responses at six hours

The model predicted that certain initial species concentrations influenced pSTAT5 levels. To determine how the initial species concentrations explain the wide variation in the six-hour responses, we analyzed the relationships between the initial species concentrations in each cell and their respective six-hour responses (**Figure 5**). The initial concentrations of both PPX and PPN seemed to be paired with six-hour *ratio*_A_ (**Figure 5A**). When either PPX or PPN was initially high in a cell, the cell would be restricted to low *ratio*_A_ at six hours. Conversely, if PPX or PPN was initially low in a cell, the cell had the opportunity to achieve high values of *ratio*_A_ at six hours. This relationship between the phosphatase concentrations and *ratio*_A_ was more pronounced with PPN than it was with PPX. In contrast, the initial concentrations of RJ and SHP2 appeared to have only a slight relationship with *ratio*_A_ (**Figure 5B**). These relationships observed for *ratio*_A_ were identical for *ratio*_B_. The initial concentration PPN seemed to be paired with six-hour *relative*_A_ (**Figure 5C**). Similar to PPN’s effect on *ratio*_A_, high PPN restricted the cell to low values for *relative*_A_ at the six-hour timepoint. RJ, SHP2, and PPX appeared to have little relationship with *relative*_A_ at six hours. These relationships observed for *relative*_A_ were identical for *relative*_B_ (**Figure 5D**).

**Figure 5.**
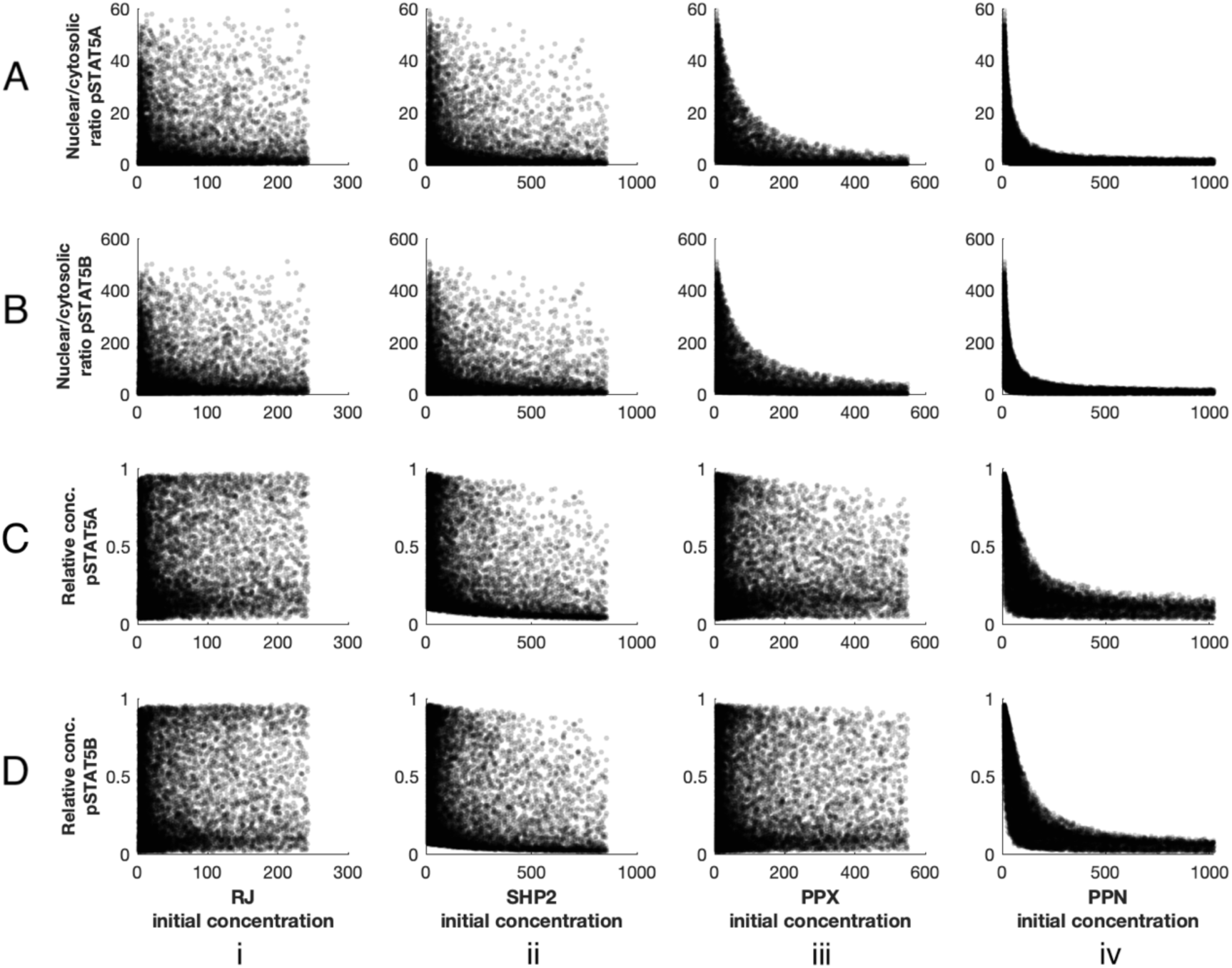
Pairwise comparisons between cell responses and initial species’ concentrations. Six-hour response values plotted against species’ concentrations for each respective cell. Values of each response are plotted along the y-axis, while concentrations are plotted along the x-axis. Each translucent dot represents one of the 10,000 simulated cells. For each cell, the (A) *ratio*_A_, (B) *ratio*_B_, (C) *relative*_A_, and (D) *relative*_B_ is plotted against its initial concentration of (i) RJ, (ii) SHP2, (iii) PPX, and (iv) PPN.

PLSR models confirmed the correlations found between the initial species concentrations and the six-hour responses. We created four PLSR models: one for each of the four responses. Each PLSR model used the initial species concentrations as inputs, and one of the four six-hour responses as the output. The models were trained on a random sample of 9,000 cells and tested on the remaining 1,000. For all four responses, the best models included two components, accounting for 93.50%, 96.28%, 91.35%, and 94.42% of the variance in the output data for *ratio*_A_, *ratio*_B_, *relative*_A_, and *relative*_B_ models, respectively.

When testing the PLSR models on the test data, each model was able to sufficiently capture the variation in the six-hour responses (**Figure 6**, left column). The R^2^ values between the PLSR model response predictions and the mechanistic model responses was greater than 0.9 for all four responses. PLSR modeling better captured the variation in responses for *ratio*_B_ (R^2^ = 0.9613) than for *ratio*_A_ (R^2^ = 0.9333). Similarly, PLSR modeling better captured the variation in responses for *relative*_B_ (R^2^ = 0.9455) than for *relative*_A_ (R^2^ = 0.9121). In addition, PLSR analysis captured the cytosolic/nuclear ratios of pSTAT5, better than the relative concentrations of pSTAT5.

**Figure 6.**
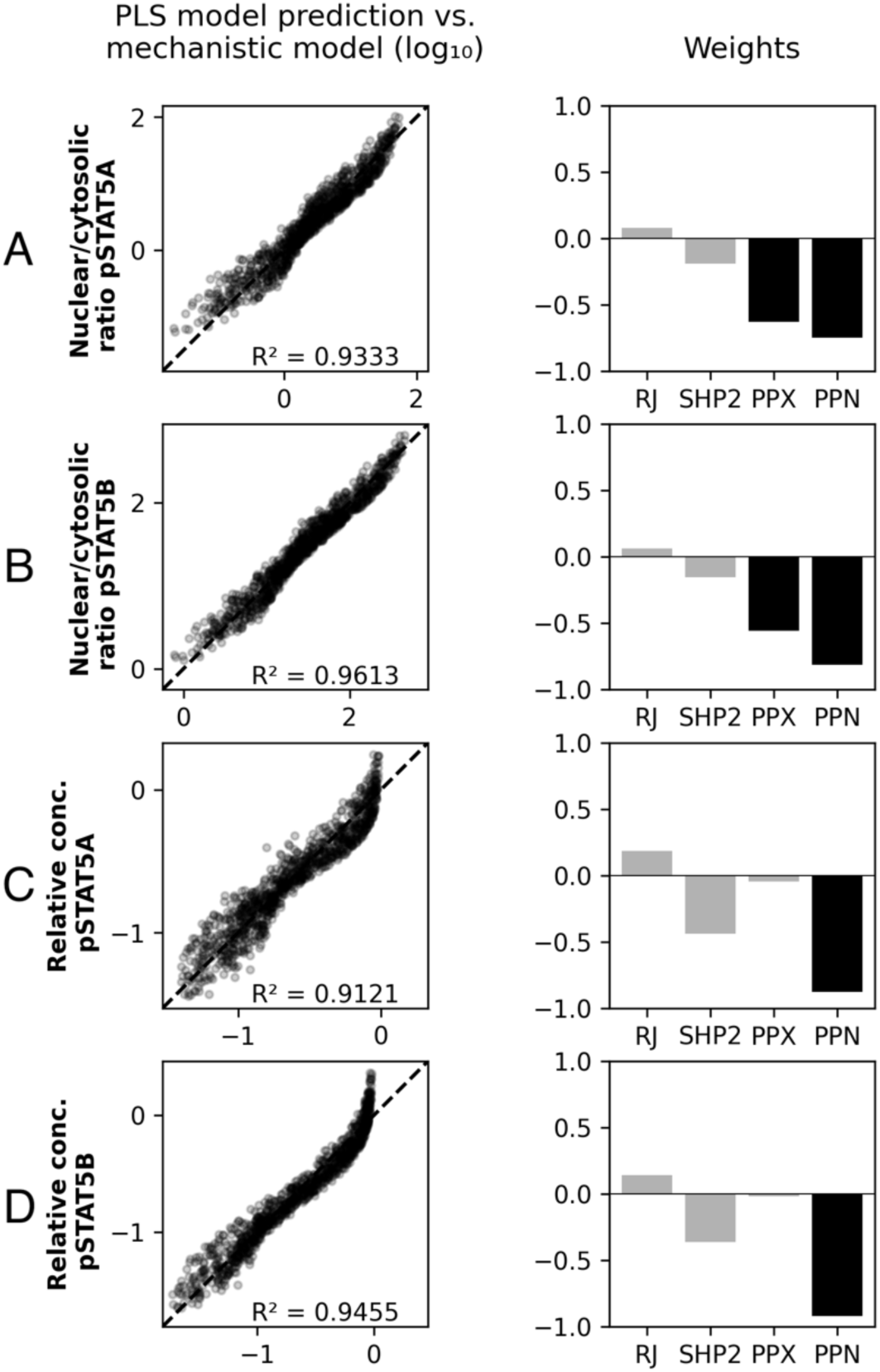
Partial least squares analysis to relate cell responses and initial species’ concentrations. PLSR model validation and input analysis for (A) *ratio*_A_, (B) *ratio*_B_, (C) *relative*_A_, and(D) *relative*_B_. (*left*) Comparison between PLSR model prediction and mechanistic model output of cell response. Log value of PLSR model cell response plotted along y-axis and log value of mechanistic model cell response plotted along x-axis. 1000 cells of testing data (not used in training of the PLSR model) plotted, with each dot representing one cell. (*right*) PLSR model weights of each initial species concentration input. Inputs with VIP score greater than or equal to one are shown with darker bars.

Using the PLSR models, we were able to identify which of the inputs most strongly influenced the output responses (**Figure 6**, right column). According to VIP scores produced by the PLSR analysis, PPX and PPN were very influential in producing the six-hour values of *ratio*_A_ and *ratio*_B_, while just PPN was shown to be influential in producing the six-hour values of both *relative*_A_ and *relative*_B_, confirming our qualitative analysis. According to the weights of the inputs, for all four response definitions, PPX and PPN had a negative influence on the six-hour responses. This means than an increase in the initial concentration of PPX or PPN decreased the pSTAT5 responses at six hours. RJ had a positive influence on all four response definitions, and SHP2 had a negative influence on all four response definitions; however, neither RJ nor SHP2 was determined to be highly influential for any of the four responses.

### Effects of initial protein concentrations on features of dynamic signaling response

In addition to analyzing the responses at one point in time, we analyzed the full six-hour time course in order to gain a deeper understanding of the overall behavior of the cell response. To do this, we identified six features of the time course response that captured the important characteristics of the peak-minimum-recovery shape: the height of the peak, the height of the minimum, the time of the peak, the time of the minimum, the absolute value of the slope from the peak to the minimum, and the slope from the minimum to the six-hour time point. As described in the Methods, **Figure 2** shows a schematic of the time course features on a sample time course. These six features were extracted from the time course for each of the four responses, creating four sets of the six features. For this analysis, we moved forward only with the cells whose response curves followed the peak-minimum recovery shape for all four responses. Therefore, we analyzed only 9,569 cells of the simulated 10,00 cells. We chose to include only these cells in our analysis since the peak-minimum-recovery shape recapitulates the dynamic response behavior that is seen in experimental observations.

The features of the overall time course for the responses varied greatly among the population of cells (**Figure 7**). The distributions of the height of the peak and height of the minimum for *ratio*_A_ were both approximately lognormal, while distributions for these features skewed left for *relative*_A_. The time of the peak had bimodal distributions for both *ratio*_A_ and *relative*_A_. The distributions of the slope features for *ratio*_A_ were approximately lognormal, while distributions for these features skewed left for *relative*_A_. The feature distributions of *ratio*_A_ had near identical shapes to the feature distributions of *ratio*_B_, and the same was true between *relative*_A_ and *relative*_B_ (**Figure S2**).

**Figure 7.**
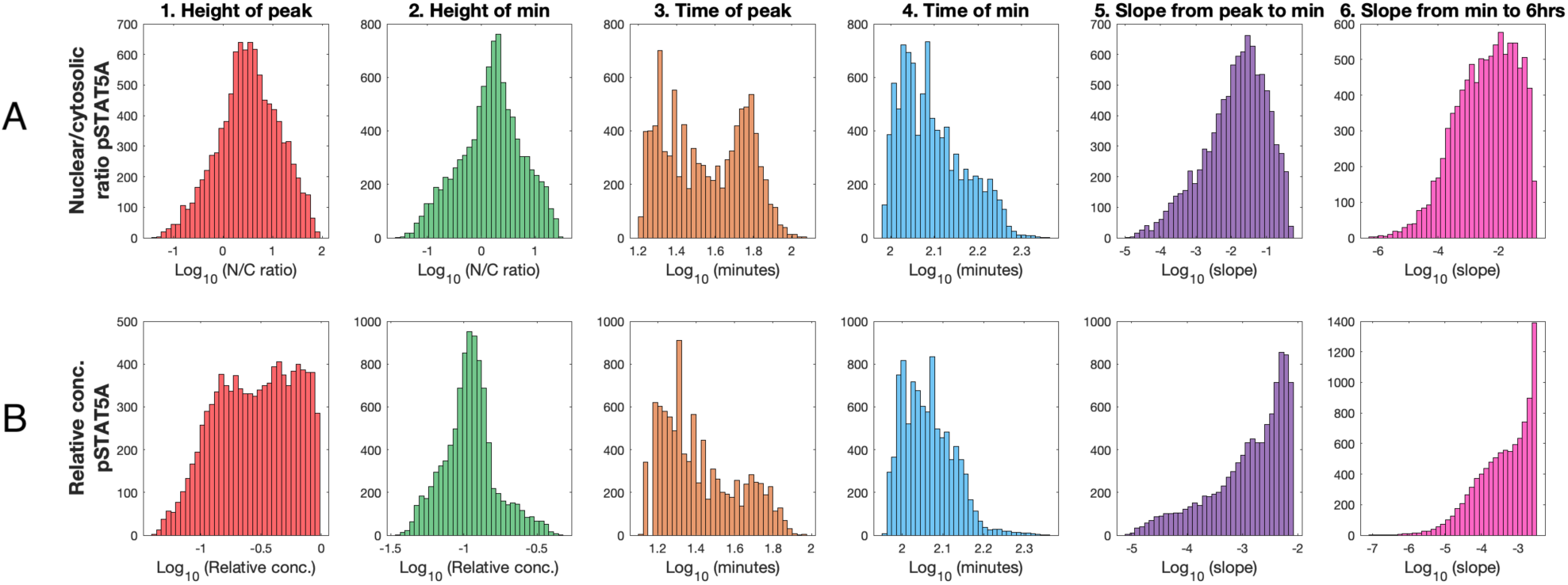
Distributions of time course features. Log-distributions of the time course response features for (A) *ratio*_A_ and (B) *relative*_A_. 9,569 data points shown. The six response features shown are: (1) height of peak, (2) height of minimum, (3) time of peak, (4) time of minimum, (5) absolute value of slope from peak to minimum, and (6) slope from minimum to six hours.

The simulations indicate which species were correlated to the six features from the full time course. Similar to our analysis of the responses at the six-hour time point, we also sought to relate the features of the full time course to the initial species’ concentrations (**Figure S3**). In this way, we could determine whether the initial species’ concentrations explain the wide variation in the time course features. For *ratio*_A_, both PPX and PPN were paired with the height of the peak, as well as with the height of the minimum. PPN appeared to be paired with the slope from the peak to the minimum, as well as with the slope from the minimum to six hours for *ratio*_A_. The pairwise relationships between the time course features and initial species concentrations appeared nearly identical between *ratio*_A_ and *ratio*_B_ (**Figure S4**), and the same was true between *relative*_*A*_ and *relative*_*B*_ (**Figures S5 and S6**). For *relative*_A_ and *relative*_B_, RJ appeared to have a relationship with the slope from the peak to the minimum, and PPN appeared to have a relationship with the slope from the minimum to six hours.

PLSR models confirmed the correlations found between the initial species concentrations and the time course response features. We created one PLSR model for each of the six features of the full time course for each of the four responses, totaling 24 models. The models were trained on 90% of the data and tested on the remaining 10%. We present the PLSR models for *ratio*_A_ in **Figure 8** and for the other three responses in **Figures S7**. Most of the PLSR models for the *ratio*_A_ response were able to sufficiently capture the variation in the time course features (**Figure 8**, left column); however, some did not accurately account for the relationship between initial concentrations and features of the time courses. The PLSR models poorly predicted the time of the peak for *ratio*_A_ (R^2^ = 0.7546) and the time of minimum for *ratio*_A_ (R^2^ = 0.3193). The inability of the PLSR models to capture the features that relate to time reveals that a linear model is not sufficient in capturing the complexity of the temporal behavior of the mechanistic model.

**Figure 8.**
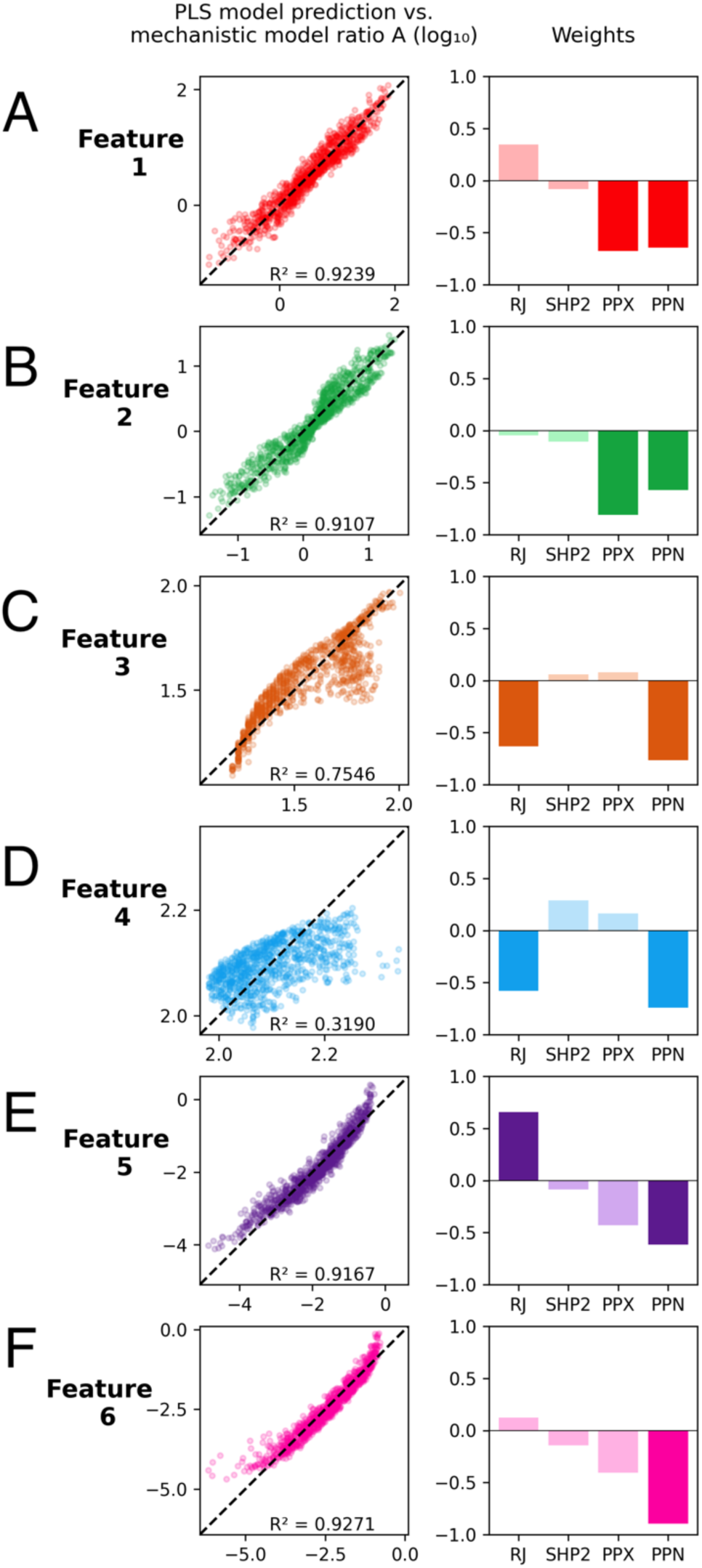
Partial least squares analysis between time course features and initial species concentrations. PLSR model validation and input analysis for six features from the *ratio*_A_ response time course: (A) height of peak, (B) height of minimum, (C) time of peak, (D) time of minimum, (E) absolute value of slope from peak to minimum, and (F) slope from minimum to six hours. (*left*) Comparison between PLSR model prediction and mechanistic model output of *ratio*_A_ time course features. Log value of PLSR model predicted features plotted along y-axis and log value of mechanistic model features plotted along x-axis. There are 957 cells of testing data (not used in training of the PLSR model) shown, with each dot representing one cell. (*right*) PLSR model weights of each initial species concentration input. Inputs with VIP score greater than or equal to one are shown with darker bars.

The VIP scores (**Figure 8**, right column) and the weights indicate that the height of the peak and height of the minimum for *ratio*_A_ were negatively influenced by PPN and PPX, confirming our qualitative findings. The VIP scores showed that the slope from the peak to the minimum for *ratio*_A_ was positively influenced by RJ and negatively influenced by PPN, and the slope from the minimum to six hours was negatively influenced by PPN.

We also considered the effect of the initial protein concentrations on the other responses of STAT5 (**Figure S7**). Between *ratio*_A_ and *ratio*_B_, the species that were determined to be significant and the directions in which the species influenced the time course features were identical. Similar to *ratio*_A_, the time of the minimum for *relative*_A_ was poorly predicted by the PLSR model (R^2^ = 0.2235). However, unlike *ratio*_A_, the time of the peak for *relative*_A_ was fairly well predicted by the PLSR model (R^2^ = 0.8500), with the model predicting that increased RJ made the peak appear earlier in the time course. In *relative*_A_, RJ had a positive influence on the height of the peak, while PPN and SHP2 had negative influences on the height of the minimum. Similar to *ratio*_A_, in *relative*_A_, RJ positively influenced the slope from the peak to the minimum, and PPN negatively influenced the slope from the minimum to six hours. Between *relative*_A_ and *relative*_B_, the species that were determined to be significant and the directions in which the species influenced the time course features were nearly identical.

## Discussion

In this work, we analyzed how heterogeneous expression of proteins influence the JAK-STAT signaling pathway in pancreatic beta cells. We developed data-driven models that can predict the outcome of this complex signaling pathway given the starting conditions of a cell. We used these models to identify relationships between the concentrations of the species in the pathway and various aspects of the cell response, which are not obvious from the connectivity of the signaling pathway. We analyzed how the species in the pathway influence the six-hour timepoint of the signaling responses, as well as the characteristics of the full time course of the signaling responses. By identifying these relationships, our analysis revealed biological targets that can potentially be used to modulate the response of pancreatic beta cells to prolactin stimulation.

We found that nuclear and cytosolic phosphatase are significant in decreasing the STAT5 response of the cell. This finding agrees with previous computational research of the JAK-STAT signaling pathway which found that nuclear phosphatase plays an important role in the JAK-STAT signaling pathway, while other species, such as the receptor and SHP2 play less important roles [24]. This information is relevant to researchers who may want to increase proliferation of beta cells via the JAK-STAT pathway, as they can use small molecule inhibitors or genetic approaches to decrease the concentrations of cytosolic and nuclear phosphatase in pancreatic beta cells. This finding is biologically relevant to potentially novel treatments for diabetes, as increasing beta cell mass could alleviate insulin insufficiency.

We also analyzed how the species’ initial concentrations affect specific mechanisms in the pathway by analyzing the characteristics of the time course response. We found that RJ has a significant, positive correlation to the magnitude of the slope from the peak to the minimum in the cell response, revealing that RJ has a role in strengthening the negative feedback mediated by SOCS. Previous research has highlighted the importance of SOCS negative feedback in inhibiting proliferation in beta cells [25] and other cell types, such as liver cells [26]. Thus, finding strategies to attenuate the strength of this negative feedback may be of particular interest to researchers studying pancreatic beta cell regeneration. Based on our conclusions, it may be possible to decrease the strength of the negative feedback by decreasing the expression of the prolactin receptor in pancreatic beta cells. This prediction that RJ can promote negative feedback to inhibit pSTAT5 response is not immediately intuitive just by examining the signaling network. This highlights the utility of our computational modeling, where the model captures the complex signaling dynamics involving positive and negative feedback loops. Additionally, we found that PPN has a significant, negative influence on the slope of the response from the minimum to six hours, indicating that decreasing PPN is a potential method to enhance the pathway’s positive feedback.

We acknowledge that the validity of our research relies on the accuracy of the mechanistic model. The parameters of the mechanistic model were fitted to experimental data, and our results are inextricably linked to the model’s ability to capture the experimental data. In particular, the available data only accounts for the first six hours of the cell’s STAT5 response upon prolactin stimulation, which limits our analysis. Additionally, certain steps in the mechanistic model were simplified, and a more detailed model may have been able to produce more accurate predictions. For example, mRNA transcription and protein translation of BcL-xL were treated as single-step reactions; however, these are processes that involve several reactions and machinery within the cell. Additionally, our research only accounts for heterogeneity in protein expression. However, heterogeneity also exists in the reaction rates, which we held constant for all cells. Future research can focus on also varying the rate constants in the mechanistic model.

Despite these limitations, our work identifies the relative importance of the initial levels of proteins in the JAK-STAT signaling pathway. We found that PPX and PPN diminish the cell response, RJ increases the strength of negative feedback, and PPN decreases the strength of positive feedback. These findings provide novel insight to the importance of modulating the mechanistic pathways that may enhance beta cell proliferation.

## Acknowledgments

The authors thank members of the Finley research group for helpful discussions and critical feedback. The work was partially supported by the USC Provost’s Undergraduate Research Fellowship awarded to A. Simoni.

## Notes

### Competing Interest Statement

The authors have declared no competing interest.

