## Supplementary Material for "Phosphatases are predicted to govern prolactin-mediated JAK-STAT signaling in pancreatic beta cells"

### Additional details regarding parameter estimation

***Background on Bayesian inference.*** In signal transduction pathways like JAK2-STAT5, it is often the case that the substrate and enzyme concentrations are comparable (1). Because a key condition for Michaelis-Menten kinetics is the assumption that substrate concentration is much larger than enzyme concentration, the signal transduction portion of this model is not governed by Michaelis-Menten kinetics. Instead, it employs mass-action kinetics. Mass-action kinetic model parameters are rate constants and initial species' concentrations. To estimate these parameters, we employed Bayesian inference, implemented via the Metropolis Hastings algorithm.

The objective of Bayesian inference is distinct, particularly in contrast to the objective of Frequentist maximum likelihood estimation: Bayesian inference sets out to approximate the posterior distribution, while maximum likelihood estimation sets out to maximize the likelihood function with respect to the model parameters. In other words, Bayesian inference is a distribution approximation problem, while maximum likelihood estimation is an optimization problem.

Further, the distribution approximated by Bayesian inference and the function maximized by maximum likelihood estimation are theoretically distinct. In the case of Bayesian inference, we approximate the posterior distribution, or conditional probability of the model parameters given experimental data. In the case of maximum likelihood estimation, we maximize the likelihood function, provided by the likelihood distribution, or conditional probability of the experimental data given the model parameters. The likelihood distribution is determined based on our observation model for the system. The posterior distribution is proportional to this likelihood distribution, multiplied by the 'prior' distribution, or prior probability of the parameters. This relationship is specified via Bayes Theorem:

$P(\vec{\theta}|\vec{x}) \propto P(\vec{x}|\vec{\theta}) * P(\vec{\theta})$

that is,

$$posterior \propto likelihood * prior$$

where:

$$\vec{\theta} = vector\ of\ model\ parameters$$
$$\vec{x} = vector\ of\ experimental\ data$$

It is evident from this formulation that a fundamental difference between the shape of the likelihood distribution and posterior distribution is the incorporation of the prior distribution; the prior provides the opportunity to incorporate informed bias into the parameter estimation process, perhaps formalizing model assumptions. For example, the parameter domain may be restricted to positive real numbers. The prior can also be made uninformative, such that the posterior distribution is proportional to the likelihood distribution.

Because of the distinct objective and distinct distribution of each procedure, the form of each result differs. On one hand, Bayesian inference generates samples of the posterior distribution. On the other hand, the maximum likelihood estimation finds a single 'best', parameterization of the likelihood distribution.

***Details regarding convergence criteria.*** The G-R method monitors convergence by comparing the variance between difference sample chains and the variance of the pooled sample chains.

Chains are initialized at different locations of the posterior; thus, the variance between sample chains is larger than the variance of the pooled chains. However, as the chains sample the same posterior distribution, the variance between sample chains approaches the variance of the pooled sample chains. Quantitatively, this means that as the chains converge, the ratio of these variances (the G-R metric) approaches one.

For the G-R method to be applicable, two key assumptions must be met. First, the chains must be initialized from a distribution over-dispersed with respect to the posterior distribution. To do this, we sampled our chains from the prior distribution used in Mortlock et al (2). This distribution is derived from an aggregated dataset of $k_{\text{cat}}$ parameters across cell types and intracellular cell pathways (3). Thus, because this model's posterior distribution describes one cell type and one pathway, this distribution was assumed to be over-dispersed with respect to our posterior distribution. The G-R method also assumes that convergence in the first and second moments (mean and variance) implies convergence in the distribution. Gaussian distributions are completely defined by their first and second moment, thus, the G-R method is applicable for Gaussian distributions. After applying a log-transformation of our marginal posterior distributions, we utilized Q-Q plots to verify that the distributions were Gaussian. With these two assumptions met, the G-R method is an applicable metric of convergence for our Bayesian inference protocol.

**Supplementary Table 1.** Parameters defining likelihood, prior, and proposal distributions.

<table>
<tr><th>Likelihood of Experimental Data Point, j</th><th>Likelihood of Experimental Data Vector</th></tr>
<tr><td>$P(x_j|\vec{\theta}) \sim \mathcal{N}\left(\mathcal{M}(\vec{\theta})_j, \sigma^2\right)$</td><td>$P(\vec{x}|\vec{\theta}) \sim \prod_j \mathcal{N}\left(\mathcal{M}(\vec{\theta})_j, \sigma^2\right)$</td></tr>
<tr><td colspan="2">$\sigma^2 \sim Inverse\ Gamma(2, .001)$<br>$x_j = experimental\ data\ point\ j$<br>$\vec{\theta} = vector\ of\ model\ parameters$<br>$\vec{x} = vector\ of\ experimental\ data$<br>$\mathcal{M}(\vec{\theta})_j = ODE\ model\ prediction\ for\ data\ point\ j, evaluated\ with\ parameter\ vector\ \vec{\theta}$</td></tr>
<tr><th>Prior Distribution of Parameter, i</th><th>Prior Distribution of Parameter Vector</th></tr>
<tr><td>$P(\theta_i) \sim LogNormal(\log(\theta_{i0}), 2)$</td><td>$P(\vec{\theta}) \sim \prod_i LogNormal(\log(\theta_{i0}), 2)$</td></tr>
<tr><td colspan="2">$\theta_i = parameter\ i$<br>$\theta_{i0} = initial\ guess\ for\ parameter\ i$</td></tr>
<tr><th>Proposal Distribution of Parameter, i</th><th>Prior of Parameter Vector</th></tr>
<tr><td>$P(\theta_{i,n}|\theta_{i,n-1}) \sim LogNormal(\log(\theta_{i,n-1}), 0.1)$</td><td>$P(\vec{\theta}_n|\vec{\theta}_{n-1}) \sim \prod_i LogNormal(\log(\theta_{i,n-1}), 0.1)$</td></tr>
<tr><td colspan="2">$\theta_{i,n-1} = \ posterior\ sample\ of\ parameter\ i\ \ from\ Metropolis - Hastings\ iteration\ n-1$</td></tr>
</table>

#### Supplementary figures

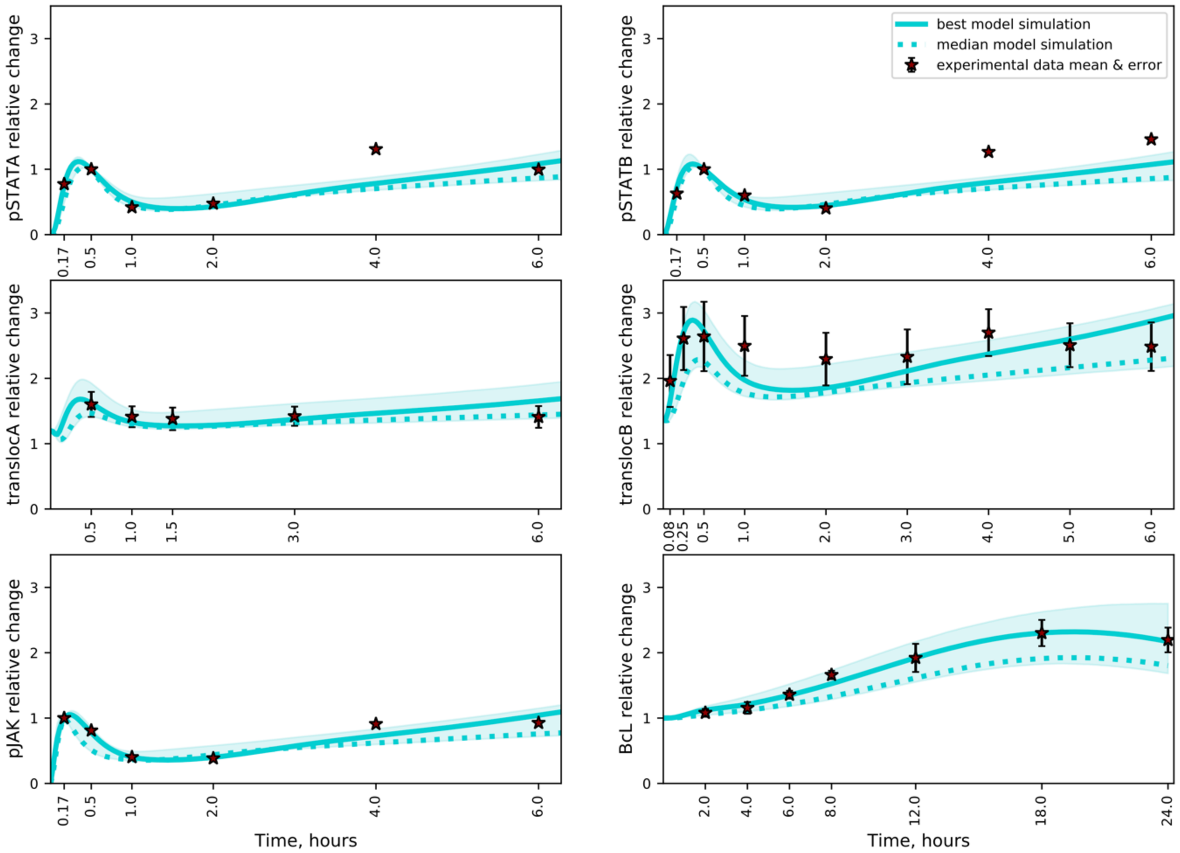


**Figure S1. Results of Bayesian parameter estimation.** Predicted signaling responses based on sampling the posterior distribution from the Bayesian parameter estimation approach. *Solid line*, best model simulation; *dashed line*, median model prediction; *shading*, the 5$^{th}$ to 95$^{th}$ quantiles of the posterior.

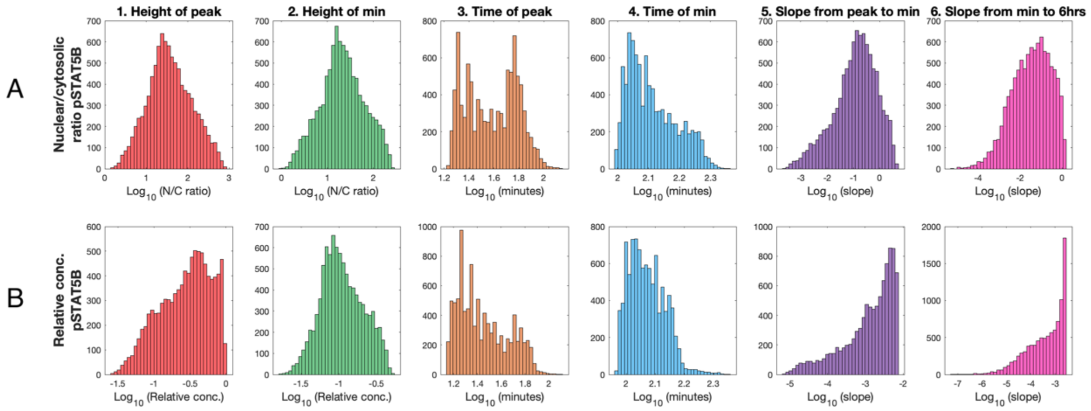


**Figure S2. Distributions of time course features.** The log-distributions of the time course response features for (A) $ratio_B$ and (B) $relative_B$. The six response features shown are: (1) height of peak, (2) height of minimum, (3) time of peak, (4) time of minimum, (5) absolute value of slope from peak to minimum, and (6) slope from minimum to six hours.

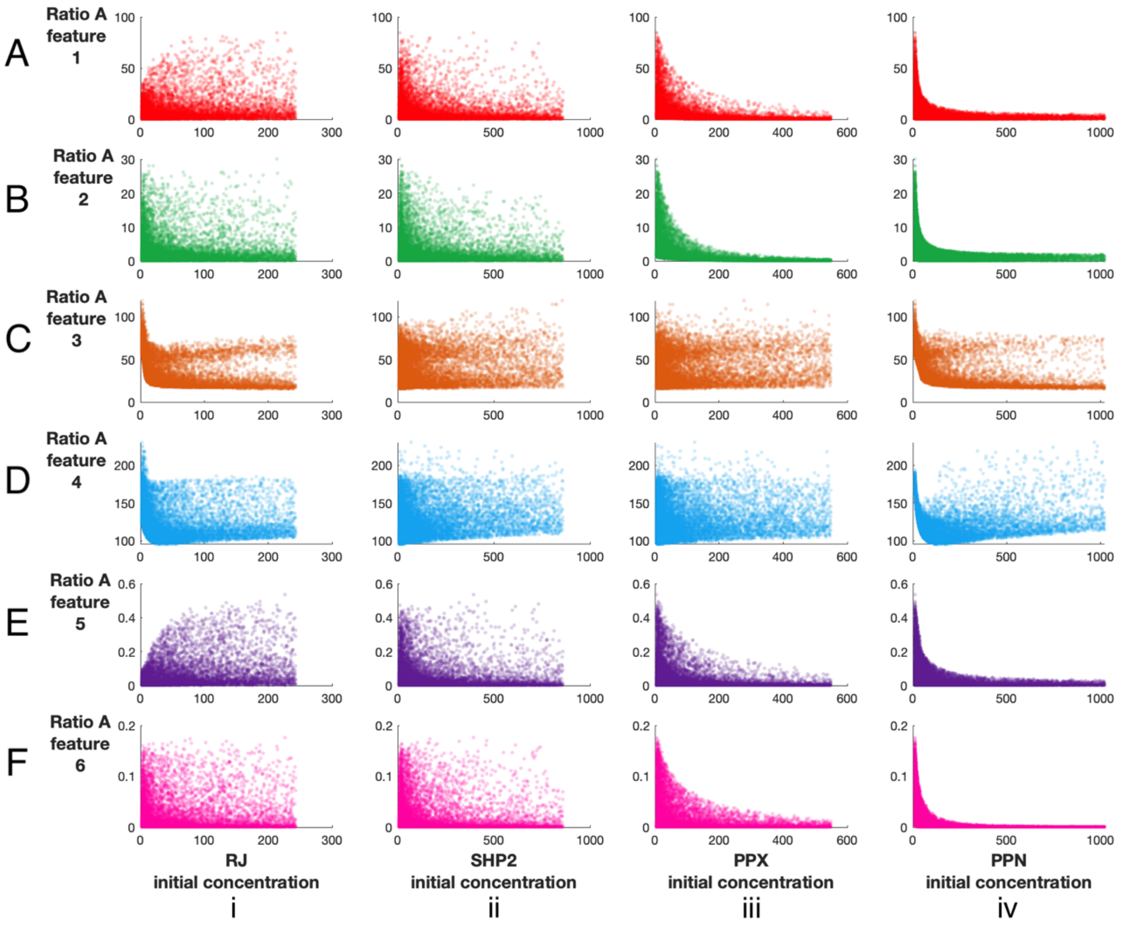


**Figure S3. Pairwise comparisons between time course features and initial species' concentrations for the *ratio*$_A$ response.** The six features of the time course for *ratio*$_A$ are plotted against the initial species' concentrations. The feature values are plotted along the y-axis, and the initial concentrations are plotted along the x-axis. The six response features shown are: (A) height of peak, (B) height of minimum, (C) time of peak, (D) time of minimum, (E) absolute value of slope from peak to minimum, and (F) slope from minimum to six hours plotted against the initial concentrations of (i) RJ, (ii) SHP2, (iii) PPX, and (iv) PPN.

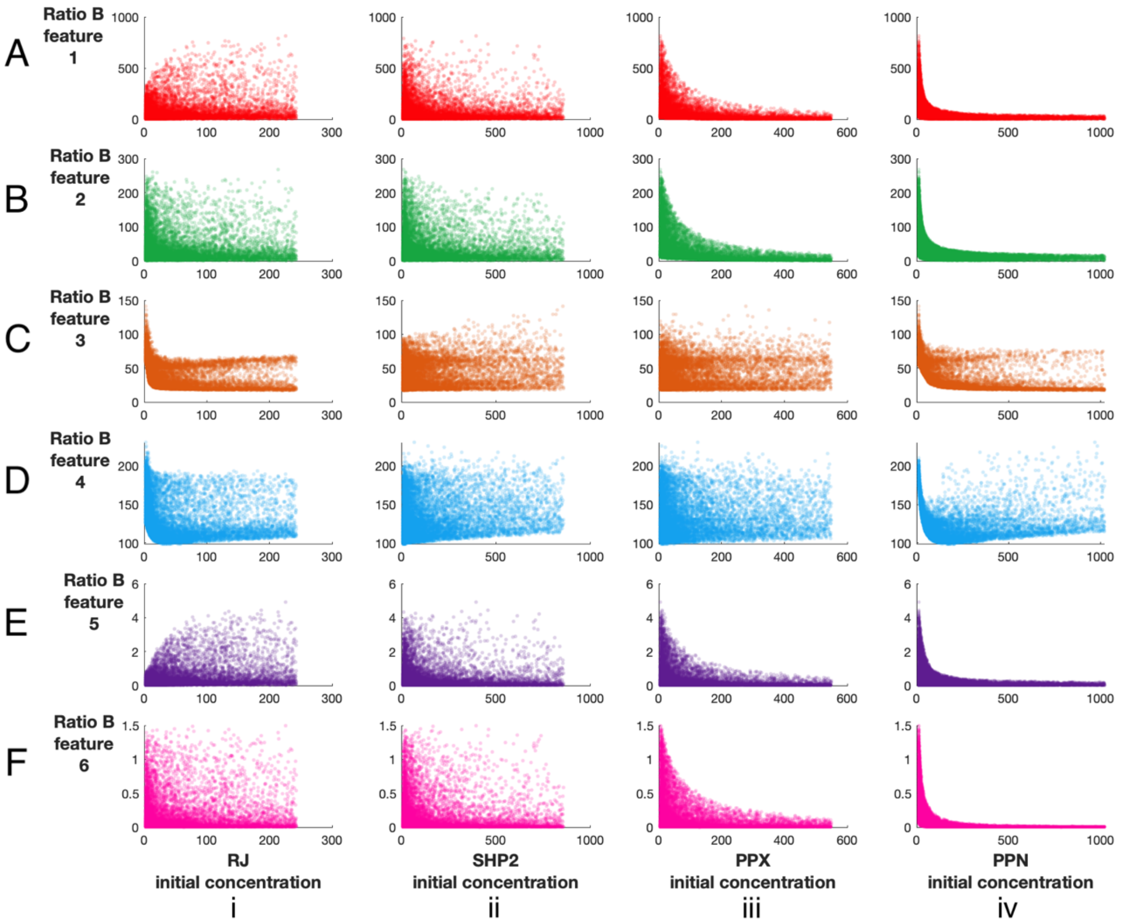


**Figure S4. Pairwise comparisons between time course features and initial species' concentrations for the *ratio*$_B$ response.** The six features of the time course for *ratio*$_B$ response are plotted against the initial species' concentrations. The feature values are plotted along the y-axis, and the initial concentrations are plotted along the x-axis. The six response features shown are: (A) height of peak, (B) height of minimum, (C) time of peak, (D) time of minimum, (E) absolute value of slope from peak to minimum, and (F) slope from minimum to six hours plotted against the initial concentrations of (i) RJ, (ii) SHP2, (iii) PPX, and (iv) PPN.

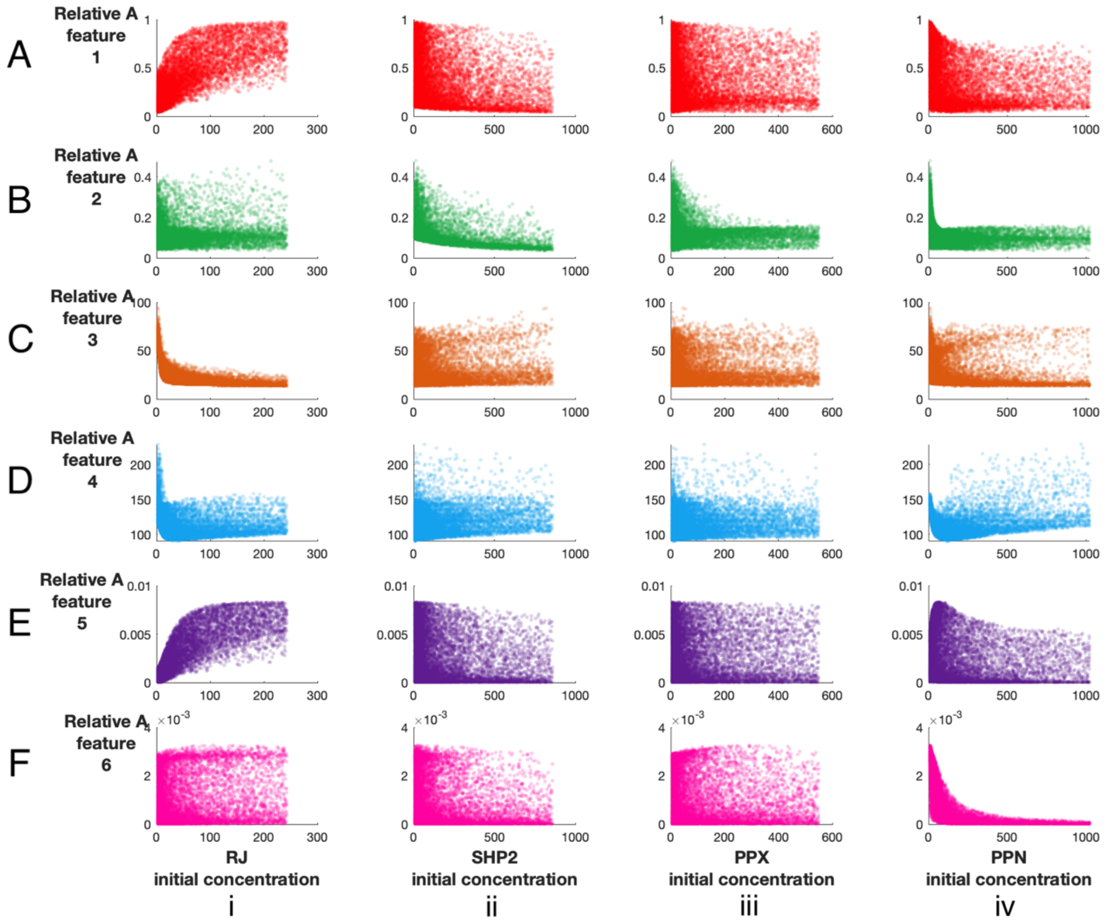


**Figure S5. Pairwise comparisons between time course features and initial species' concentrations for the *relative*$_A$ response.** The six features of the time course for the *relative*$_A$ response are plotted against the initial species' concentrations. The feature values are plotted along the y-axis, and the initial concentrations are plotted along the x-axis. The six response features shown are: (A) height of peak, (B) height of minimum, (C) time of peak, (D) time of minimum, (E) absolute value of slope from peak to minimum, and (F) slope from minimum to six hours plotted against the initial concentrations of (i) RJ, (ii) SHP2, (iii) PPX, and (iv) PPN.

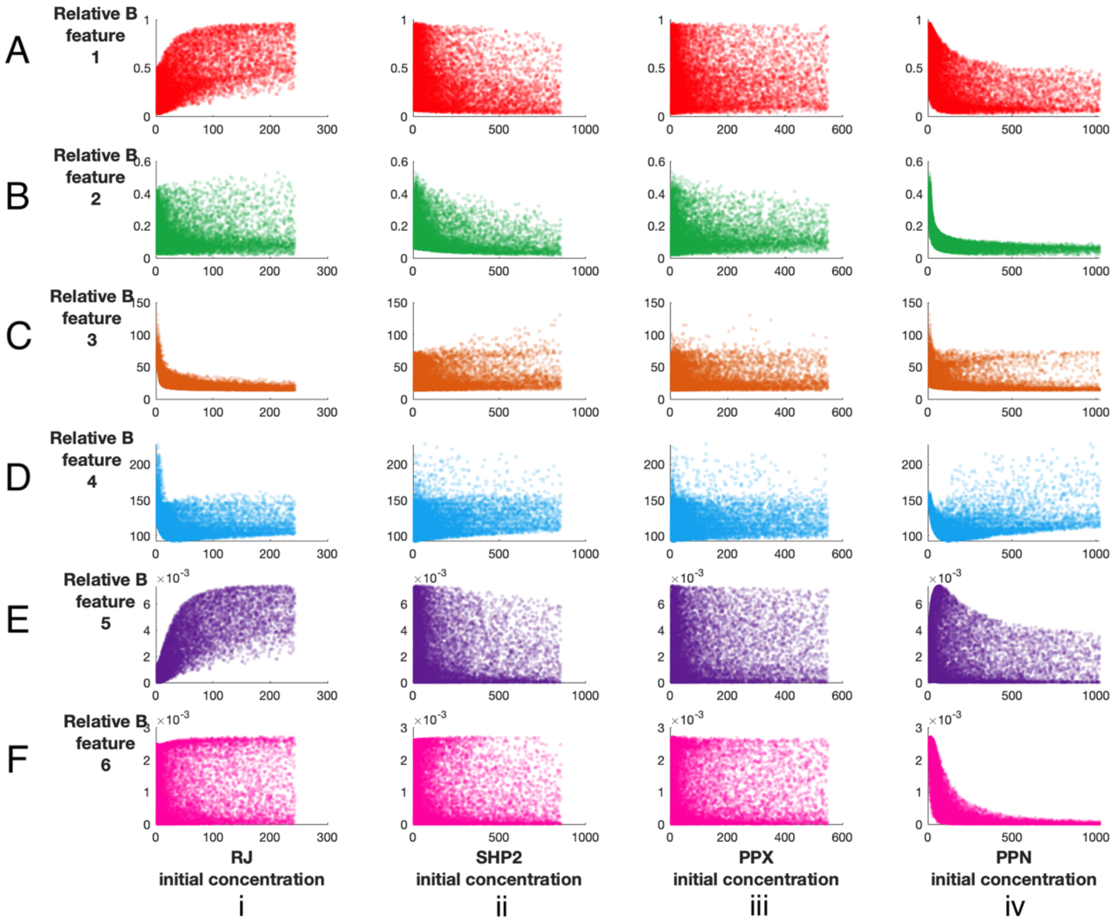


**Figure S6. Pairwise comparisons between time course features and initial species' concentrations.** The six features of the time course for the *relative*$_B$ response are plotted against the initial species' concentrations. The feature values are plotted along the y-axis, and the initial concentrations are plotted along the x-axis. The six response features shown are: (A) height of peak, (B) height of minimum, (C) time of peak, (D) time of minimum, (E) absolute value of slope from peak to minimum, and (F) slope from minimum to six hours plotted against the initial concentrations of (i) RJ, (ii) SHP2, (iii) PPX, and (iv) PPN.

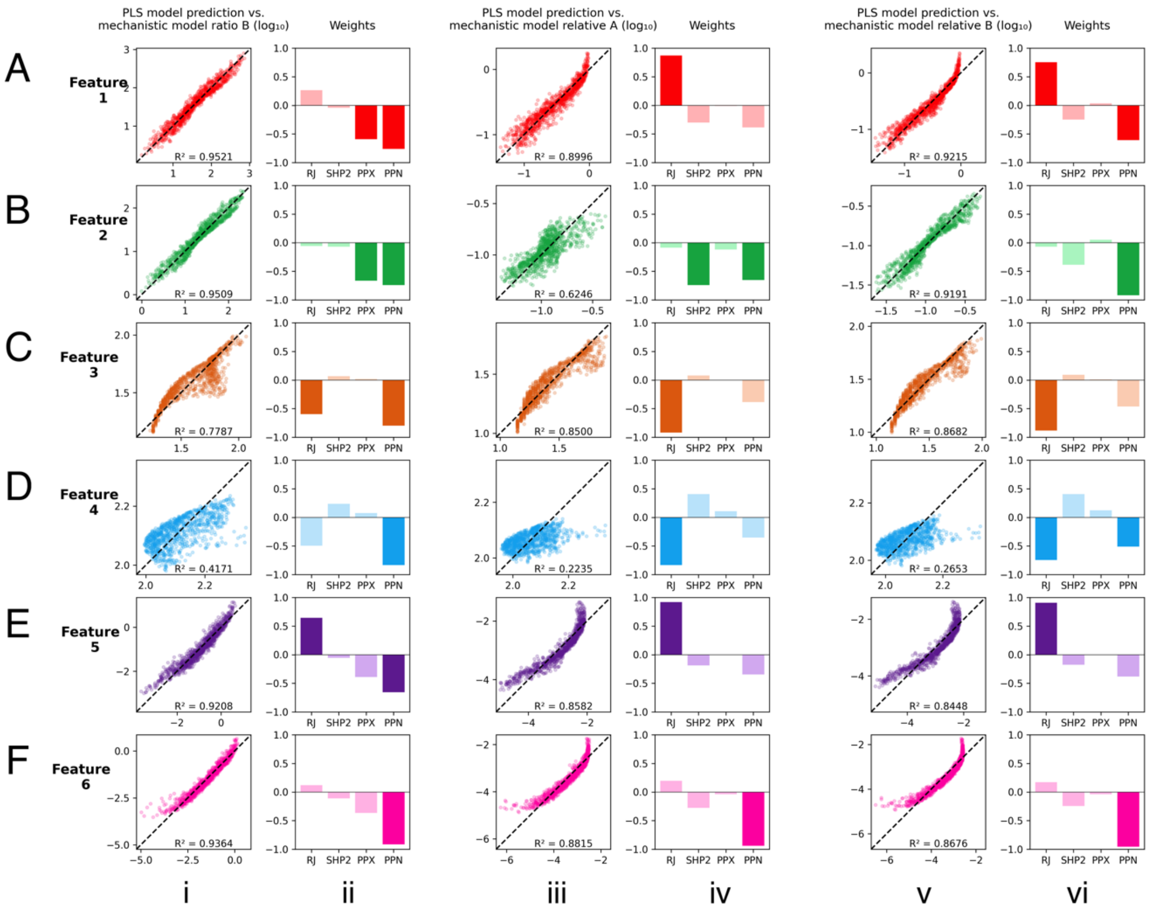


**Figure S7. Partial least squares analysis between time course features and initial species' concentrations.** PLSR model validation and input analysis for six features: (A) height of peak, (B) height of minimum, (C) time of peak, (D) time of minimum, (E) absolute value of slope from peak to minimum, and (F) slope from minimum to six hours. (i) PLSR model validation and (ii) PLSR model weights VIP scores for $ratio_B$ response, (iii) PLSR model validation and (iv) PLSR model weights VIP scores for $relative_A$ response; (v) PLSR model validation and (vi) PLSR model weights VIP scores for $relative_B$ response.
